# *In Vitro* Assay for the Detection of Network Connectivity in Embryonic Stem Cell-Derived Cultures

**DOI:** 10.1101/377689

**Authors:** Jeffrey R Gamble, Eric T Zhang, Nisha Iyer, Shelly Sakiyama-Elbert, Dennis L Barbour

## Abstract

Stem cell transplantation holds great promise as a repair strategy following spinal cord injury. Embryonic stem cell (ESC) transplantation therapies have elicited encouraging though limited improvement in motor and sensory function with the use of heterogeneous mixtures of spinal cord neural progenitors and ESCs. Recently, transgenic lines of ESCs have been developed to allow for purification of specific candidate populations prior to transplantation, but the functional network connectivity of these populations and its relationship to recovery is difficult to examine with current technological limitations. In this study, we combine an ESC differentiation protocol, multi-electrode arrays (MEAs), and previously developed neuronal connectivity detection algorithms to develop an *in vitro* high-throughput assay of network connectivity in ESC-derived populations of neurons. Neuronal aggregation results in more consistent detection of individual neuronal activity than dissociated cultures. Both aggregated and dissociated culture types exhibited synchronized bursting behaviors at days 17 and 18 on MEAs, and thousands of statistically significance functional connections were detected in both culture types. Aggregate cultures, however, demonstrate a tight linear relationship between the inter-neuron distance of neuronal pairs and the time delay of the neuronal pair functional connections, whereas dissociated cultures do not. These results suggest that ESC-derived aggregated cultures may reflect some of the spatiotemporal connectivity characteristics of *in vivo* tissue and prove to be useful models of investigating potentially therapeutic populations of ESC-derived neurons *in vitro*.

**NOVELTY AND SIGNIFICANCE:** Previous investigations of stem cell-derived network connectivity on multi-electrode arrays (MEAs) have been limited to characterizations of bursting activity or broad averages of overall temporal network correlations, both of which overlook neuronal level interactions. The use of spike-sorting and short-time cross-correlation histograms along with statistical techniques developed specifically for MEAs allows for the characterization of functional connections between individual stem cell-derived neurons. This high-throughput connectivity assay will open doors for future examinations of the differences in functional network formation between various candidate stem cell-derived populations for spinal cord injury transplantation therapies—a critical inquiry into their therapeutic viability.

## INTRODUCTION

In recent decades stem cell therapy has become promising as a potential treatment strategy for spinal cord injury (SCI). Typically, heterogeneous mixtures of embryonic stem cells (ESCs), neural-restricted progenitors, or glial-restricted progenitors are loaded into gels or scaffolds that are then either injected or surgically inserted into the spinal cavity (Duncan, Aguayo, et al. 1981; Brustle, Jones, et al. 1999; Nori, Okada, et al. 2011; Tetzlaff, Okon, et al. 2011; Mothe, Tam, et al. 2013; McCreedy, Wilems, et al. 2014). One rationale for this intervention is that the grafted cells have the capacity to differentiate into neurons or glia and incorporate themselves into the endogenous circuitry spared by the injury, forming new spinal circuits within the grafted population and between the grafted cells and host cells. These circuits can then serve as neuronal relays for communication along the injured cord (Abematsu, Tsujimura, et al. 2010; Bonner, Connors, et al. 2011; Fujimoto, Abematsu, et al. 2012; Lu, Wang, et al. 2012; Hou, Tom, et al. 2013; Sharp, Yee, et al. 2014).

Although a spectrum of *in vitro* protocols has been developed to consistently differentiate ESCs into well-characterized populations enriched for predetermined neural subtypes and to purify the spinal cord neuronal subpopulation of choice through genetic-engineering (Li, Pevny, et al. 1998; Anderson, Self, et al. 2007; McCreedy, Rieger, et al. 2012; Brown, Butts, et al. 2014; McCreedy, Brown, et al. 2014; Xu, Iyer, et al. 2015; Xu and Sakiyama-Elbert 2015; Iyer, Huettner, et al. 2016), the intrinsic complexity of the post-SCI environment, as well as severe limitations of traditional *in vivo* technologies, currently present challenges to studying intra-graft connectivity properties post-transplantation (Bonner, Connors, et al. 2011; Lee, Lane, et al. 2014). *In vitro* electrophysiology methodologies from other fields, in conjunction with cell induction and cell culture methods developed within the SCI field, present a compelling alternate platform to investigate how different stem cell-derived subpopulations interconnect within the overall neuronal population.

In particular, functional connectivity, referring to a correlation between two time series with no assumption about the underlying circuit giving rise to the measured relatedness, has proven to be a useful framework for assessing neuronal interconnectivity when anatomical connectivity is inaccessible experimentally (Moore, Perkel, et al. 1966; Perkel, Gerstein, et al. 1967; Gerstein and Perkel 1972; Aertsen, Gerstein, et al. 1989; Aertsen, Vaadia, et al. 1991; Friston, Frith, et al. 1993; Friston 1994; Friston 2011). Due to their parallel channels of extracellular recording, planar multi-electrode arrays (MEAs) have been shown to be particularly well-suited to analyze functional connectivity for the purpose of characterizing network development, plasticity, and structure *in vitro* (Chao, Bakkum, et al. 2007; Garofalo, Nieus, et al. 2009; Maccione, Garofalo, et al. 2012; Freeman, Krock, et al. 2013; Cutts and Eglen 2014; Poli, Pastore, et al. 2015; Pastore, Poli, et al. 2016). These investigations typically measure functional connectivity by assessing individual connections and their associated properties (Maccione, Garofalo, et al. 2012; Freeman, Krock, et al. 2013) or by constructing topological maps to facilitate comparisons of network structure across conditions or groups (Maccione, Gandolfo, et al. 2010; Boehler, Leondopulos, et al. 2012; Downes, Hammond, et al. 2012; Marconi, Nieus, et al. 2012; Poli, Pastore, et al. 2015; Schroeter, Charlesworth, et al. 2015), although the mathematical validity of comparing networks with differing quantities of neuronal nodes has recently been called into question (van Wijk, Stam, et al. 2010).

Only in recent years have MEAs been established as a modality for functionally analyzing the connectivity properties within populations of cultured ESC-derived neurons (Ban, Bonifazi, et al. 2007; Illes, Fleischer, et al. 2007; Heikkila, Yla-Outinen, et al. 2009; Illes, Jakab, et al. 2014). Ban and colleagues opened the door for more in-depth inquiries into the spatial and temporal properties of ES-derived neuronal interactions by demonstrating, with the use of MEAs, the capacity for ES-derived neurons to form functional networks (Ban, Bonifazi, et al. 2007). Multiple groups have shown that this spatial activity can be modulated both in human and mouse stem cell-derived populations by the application of neurotransmitters and channel antagonists, suggesting the expression of synaptic ionic channels native to endogenous neuronal populations (Illes, Fleischer, et al. 2007; Heikkila, Yla-Outinen, et al. 2009; Lappalainen, Salomaki, et al. 2010).

However, these fundamental characterizations of ESC-derived *in vitro* neural networks have yet to assess how individual neurons functionally interconnect, an assessment likely to capture nuanced functional differences between ESC-derived populations that may actually be correlated with their efficacy, or lack thereof, as therapeutic ESC-derived spinal cord networks (Bonner and Steward 2015). Applications of such methodologies would enable investigations of fundamental questions about ESC-derived networks pertaining to their use as grafts, such as the relationship between neuron-neuron functional connectivity and cell culture methodology (i.e. dissociated vs aggregated cultures), which has previously been shown to influence how closely neuronal cultures mimic *in vivo* environments (Pampaloni, Reynaud, et al. 2007; Lu, Searle, et al. 2012; Edmondson, Broglie, et al. 2014). For example, the effects of neural aggregation on long-term function of ESC-derived culture have been explored previously at the level of neural activity, demonstrating that aggregation leads to increasingly synchronous bursting across electrodes (Illes, Theiss, et al. 2009).

In the current study we extend the previously discussed MEA investigations of ESC-derived interconnectivity by investigating the impact of cell culture methodologies on neuron-neuron functional connectivity. We do so by combining planar MEAs and a computational methodology recently shown to be capable of mapping thousands of pairwise functional connections between spike-sorted neurons in neuronal cultures (Freeman, Krock, et al. 2013). Cross correlation, the measure of connectivity utilized here, has historically been used to detect a spectrum of neuronal interconnectivity, from neuron-neuron monosynaptic connections to polysynaptic synchrony (Binder and Powers 2001; Constantinidis, Franowicz, et al. 2001; Herrmann and Gerstner 2002; Turker and Powers 2002; Bartho, Hirase, et al. 2004; Veredas, Vico, et al. 2005; Fujisawa, Amarasingham, et al. 2008; Ostojic, Brunel, et al. 2009). The assay presented here not only detected differences in neuronal activity between dissociated and aggregated cultures, but also captured in a high-throughput manner differences between culture methodologies pertaining to the spatiotemporal characteristics of neuron-neuron functional connectivity, which has implications for advancing assessments of ESC-derived neuronal for SCI repair.

## METHODS

### ESC Culture

Cultures enriched for Chx10-expressing V2a interneurons (V2a INs) were induced as previously described (Brown, Butts, et al. 2014). A mouse ESC line expressing TdTomato under the Bactin promotor was maintained in T-25 flasks in complete media, consisting of Dulbelcco’s Modified Eagle Medium (DMEM; Life Technologies #11965-092) containing 10% newborn calf serum (Life Technologies #26140-079), and 1× Embryomax Nucleosides (Millipore #ES-008-D). Every two days, ESCs were passaged at a 1:5 ratio with fresh complete media supplemented with 100 μM β-mercaptoethanol (βME; Life Technologies #21985-023) and 1000 U/mL leukemia inhibitory factor (LIF; Millipore #ESG1106).

### Chx10 induction (2^−^/4^+^)

At the beginning of the induction, 10^6^ ESCs were suspended in 10 mL of DFK5 media, which consists of a DMEM/F12 base supplemented with βME, 1:200 100× Embryomax Nucleosides, 50 μM nonessential amino acids (Life Technologies #11140-050), 100× insulin transferrin-selenium (Life Technologies #41400-045) and 5% knockout replacement serum (Life Technologies #10828-028) on an agar-coated 100 mm petri dish for two days in order to form embryoid bodies (EBs) (2^−^). Media was then aspirated from the EBs and changed to DFK5 media containing 10 nM retinoic acid (RA; Sigma #R2625) and 1 μM purmorphamine (EMD Millipore #540223) for 2 days (2^+^) prior to the media being aspirated and changed to DFK5 with 10 nM RA, 1 μM purmorphamine, and 5 μM of a gamma secretase inhibitor N-[N-(3,5-difluorophenacetyl-L-alanyl)]-(S)-phenylglycine t-butyl ester (DAPT; Sigma-Aldrich #D5942) for the last two days (4^+^).

### Dissociation and Aggregation

After the 6-day induction protocol, EBs were either dissociated or underwent a neural aggregation protocol prior to plating for MEA cultures. For dissociation-alone cultures, EBs were collected, suspended in 0.25% Tryspin-EDTA (Life Technologies #25200-056) for 10 minutes, and mechanically separated by repeated pipetting. For neural aggregation cultures, EBs were initially dissociated as described, but were then plated on poly-L-ornithine (Sigma-Aldrich #P4957) and laminin-coated T75 flasks at a density of 5×10^5^ cells/cm^2^ in DFK5 supplemented with B-27 (Life Technologies #17504-044), glutaMAX (Life Technologies #35050-061), 5 ng/mL glial-derived neurotrophic factor (GDNF; Peprotech #450-10), 5 ng/mL brain-derived neurotrophic factor (BDNF; Peprotech #450-02) and 5 ng/mL neurotrophin-3 (NT-3; Peprotech #450-03). After 24 hours, the flask cultures were washed twice with DMEM/F12 and then lifted from flasks using Accutase (Sigma, #A6964) for 30 min at room temperature. A total of 5×10^5^ cells were seeded into each well of a 400 μm Aggrewell plate (StemCell Technologies, #27845) in a modified DFKNB media consisting of DFK5 and Neurobasal media (NB; Life Technologies #21103-049) supplemented with B-27, glutaMAX, 5 ng/mL GDNF, 5 ng/mL BDNF, and 5 ng/mL NT-3 for 2 days of aggregate formation. Half the media was replaced daily. Aggregates were lifted by trituration and allowed to settle in micro-centrifuge tubes prior to plating as appropriate.

### MEA cell culture

Either dissociated cells or aggregated cells were plated on individual 8×8 electrode grid S1-type MEAs with 200 μm electrode spacing and 10 μm electrode contact diameter (Warner Instruments #890102) that had been coated with poly-L-ornithine/laminin. Dissociated cells were plated uniformly at a cell density of 5×10^5^ cells per MEA, while neural aggregates were plated at the center of each MEA at a cell density of 1.5×10^5^. All cultures were incubated at 37° C in a humidity-controlled, 5% CO_2_ environment for the entire culture period. Both types were cultured in DFKNB with B-27, glutaMAX, 5 ng/mL GDNF, 5 ng/mL BDNF, and 5 ng/mL NT-3. Full media changes of the supplemented DFKNB media were performed every two days up to day 6 in culture. After 6 days, the media was replaced with NB containing the same supplements for the duration of culture, with a full change of media performed every two days. Images of MEA cell cultures were captured viewing from beneath the MEA dish using an Olympus IX70 inverted microscope with an attached MICROfire camera.

### MEA recording

The recorded signals were amplified with a Multichannel Systems MEA2100 60-channel signal amplifier (ALA Scientific Instruments, Inc). The Multichannel Systems software gain was set to 1200× with an analog-to-digital sampling rate of 20 ksamples/s. Channels with high-amplitude noise were turned off in order to avoid the spread of noise to other channels. All experiments were conducted at 37° C with the use of a MultiChannelSytems temperature-controller (ALA Scientific Instruments, Inc) in an open-air table-top environment with no replacement of media during recording. Recordings were performed using the MultiChannel Systems data acquisition software, MC Rack, during which all 60 channels were filtered in parallel with a 4^th^ order digital Butterworth band-pass filter from 300 Hz to 5000 Hz. Neural activity of all non-noisy channels was recorded for one hour, which ensured enough spiking activity to construct meaningful cross-correlograms. For each one-hour recording, only neuronal spikes passing a 5 standard deviation negative voltage deflection threshold, along with the time stamp of the threshold crossing, were extracted and saved to disk. Each spike waveform consisted of 1 ms of samples pre-threshold crossing and 2 ms of samples post-threshold crossing.

### Offline Spike-sorting

Extracted spike waveforms were spike sorted offline by the MC Rack software, then imported into MATLAB (Mathworks, Inc). The spike time associated with each waveform initially corresponded to the time at which the negative deflections of action potentials crossed the negative thresholds. Once imported into MATLAB, the sample rate was used to shift the spike times to correspond to the minimum point in the negative deflection of the spike waveforms in order for the waveforms to be evenly aligned during feature extraction. Waveforms were also trimmed to include only 0.5 ms before the minimum value of the negative deflection and 1.5 ms after the minimum value of the negative deflection. Only identifiable single neuron spikes were included in the analysis. For feature extraction, principal components analysis (PCA) was performed on a feature matrix created by stacking the vectors corresponding to each waveform on a single channel. The output matrix corresponding to the 1–5 principal components containing the most variance were the inputs for K-means clustering, which was then used to assign centroids to a predetermined number of clusters (number of identified neurons) and index each PCA vector corresponding to a single waveform to a cluster. This analysis identified anywhere from one to five individual neurons for each channel.

### Bursting Quantification

The level of bursting in the cultures, also called “burstiness,” was quantified as described by Wagenaar and colleagues (Wagenaar, Madhavan, et al. 2005). Recordings were divided into 1-second bins. The number of spikes across all individual neurons was counted for each 1-second bin (summed across the network). Examining the 15% of bins with the largest number of spike counts, the fraction of spikes across the entire recording period accounted for by these top 15% bins was then computed and labeled f_15_. A f_15_ close to 0.15 implies evenly distributed firing temporally, or tonic firing, across the network. Highly synchronized bursting across the network followed by periods of silence will result in a f_15_ close to 1. The burstiness index BI is then computed as

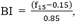

where BI equal to 0 implies tonic firing and BI equal to 1 implies dominating bursting.

### Spike-timing cross-correlation histogram (CCH)

An exact replication of the spike-timing cross-correlation histogram (CCH) procedure utilized in Freeman, *et al*., was used as the measure of functional connectivity in this study (Fujisawa, Amarasingham, et al. 2008; Freeman, Krock, et al. 2013). Only neurons containing at least 100 identified action potentials over the hour-long recording window were included in the CCH analysis. For the first spike-time in neuron A’s spike train, a 2005 ms window was centered around the spike-time such that the location in time at which the spiked occurred became 0 on the spike delay axis. The window was then divided into 5 ms bins by centering a bin at time 0 on the delay axis and shifting the 5 ms bin in 1 ms increments in both the positive and negative directions. With this current spike from train A as a frame of reference at delay 0, the number of spikes in train B that fell into each spike delay bin were counted, leaving a histogram of the spike delay counts for a single spike in train A. These operations were then repeated for each spike in train A. The spike-timing CCH for neuron A compared to neuron B was then generated by summing all of the single spike histograms for neuron A and normalizing by the square root of the product of the total spike counts for neurons A and B across the entire window. We then low-pass filtered the final result with a 5-point boxcar filter.

Comparison of connections relied on the calculation of z-scores (Freeman, Krock, et al. 2013). Z-scores are indicators of whether a neuron modulated the spiking of a target neuron and was calculated as follows:

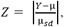

where *Y* is the value at the peak/trough of the maximum deflection closest to zero, μ is the mean of the particular cross-correlogram, and μ_sd_ is the standard deviation of μ for all crosscorrelograms.

### Significance Calculation of Functional Connections

In order to determine which neuronal connections were statistically significant, we used the Between-Sample Analysis of Connectivity (BSAC) methodology described in Freeman, *et al*., to differentiate between spurious and meaningful connections (Freeman, Krock, et al. 2013). For all MEA cultures used within a single experimental condition, cross-correlograms and corresponding z-scores were calculated for every possible neuron pair consisting of isolated neurons existing in separate, physically distinct MEAs. Because the comparison in this control condition is between two neurons existing in two different dishes, any correlation between their activities is not physiologically relevant. Using this paradigm, we built distributions of inter-culture z-scores (functional connections between two neurons located in different MEA dishes) for positive connections that could be used to empirically determine a z-score threshold for statistical significance in the intra-culture distribution (functional connections between neurons located within the same MEA dish), circumventing having to make any assumptions about spike train characteristics that are typically necessary to compute functional connection significance. We chose to use the z-score threshold corresponding to a false-discovery rate (FDR) of 0.05, meaning 5% of the inter-culture connection z-score data lay above the derived threshold value.

This positive connection threshold was then used on a z-score distribution constructed from intra-culture comparisons. For the same MEAs used for the inter-culture analysis, cross-correlations and corresponding z-scores were calculated for every possible neuron pair consisting of spike-sorted neurons existing within the same MEA. These putative connections had the potential to be physiologically relevant. A positive connection distribution was again constructed, this time from within-culture comparisons. The threshold determined from the across-culture distribution was then applied to the positive intra-culture distribution. Any connection with a z-score lying at the 5% FDR threshold or better was deemed significant and used in further analyses.

## RESULTS

### Growth of ES-Derived Chx10-enriched MEA Cultures

ES-derived neural populations were induced towards V2a interneurons that are marked with increased expression of Chx10 using the six day 2^−^/4^+^ protocol. Chx10-enriched populations were grown on MEAs in two different manners: dissociated or aggregated cultures (**Figure 1**). Dissociated Chx10-enriched populations formed a fairly uniform layer of both neurons and glia across the MEAs. For all of the dissociated cultures (n = 6), the majority of the 60 electrodes in each culture had cell bodies in close proximity to or on top of the electrode contacts shortly after seeding (**Fig1a-b**), but a subset of cultures exhibited some recession of growth that resulted in a reduction of electrode coverage. Dissociated cultures remained healthy up to the recording time points, either DIV 17 or 18 as measured by the persistent spontaneous activity in the cultures. Neural aggregates of the ES-derived Chx10-enriched cells were plated on the MEAs such that the majority of the aggregates sat in the middle of the MEA dish in order to ensure coverage of the electrode contacts with neurons. Maturing aggregate cultures formed an extremely dense layer over the electrode contacts with some migration of glia radially from the main body of aggregates (**Fig1c-d**). Aggregate cultures (n = 4) also remained healthy up to the electrophysiological recordings at DIV 17 or 18 as measured by the persistent spontaneous activity in the cultures.

**Figured 1.**
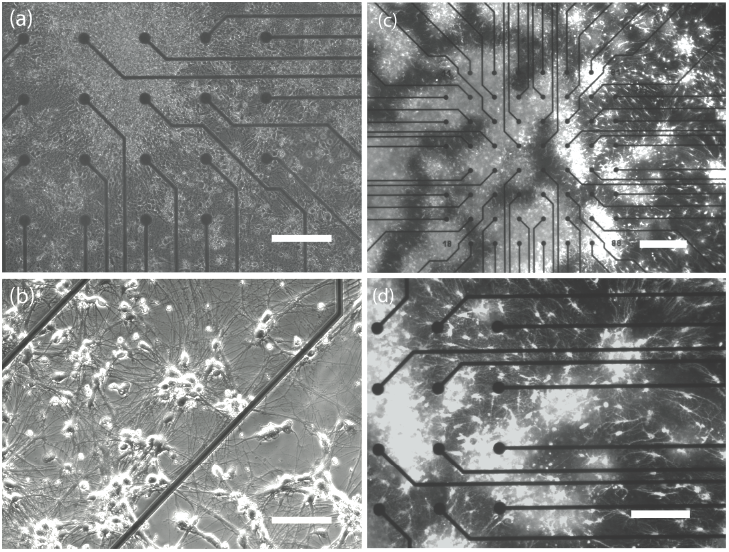
Cultures of ES-derived Chx10-enriched populations of neurons on MEAs. **(a)** 10× phase image of ES-derived dissociated neuronal cultures densely growing over the electrode contacts (black circles) at 14 days in MEA culture **(b)** 20× phase image of ES-derived dissociated neuronal cultures growing in small clusters on a MEA. **(c)** 4× fluorescent image of ES-derived aggregated neuronal cultures densely growing over the electrode contacts (black circles) at 14 days in MEA culture. All ESC-derived cells are fluorescently labeled. **(d)** 10× fluorescent image of ES-derived neural aggregates clustered over the MEA electrode contacts. (Scale bars = 200 μm, 100 μm, 400 μm, and 200 μm respectively)

### MEAs Recorded Spontaneous Synchronized Activity

MEAs recorded spontaneous network activity in both dissociated and aggregated cultures at either DIV 17 or 18. Both culture types exhibited bursting activity synchronized across the MEA, with the synchronized bursts occurring in two groupings of firing closely positioned in time followed by long periods of silence (**Figure 2a**). Closer inspection of individual spike clusters showed variable levels of spike density within the bursts. **Figure 2b** is an exemplar of a dense burst from an electrode in a dissociated culture, while **Figure 2c** is an increasingly sparse burst from the same culture. Of the 6 dissociated MEA cultures and the 4 aggregated MEA cultures, all cultures displayed double bursting behavior, which is consistent with previous investigations of MEA activity in embryonic-derived cultures between 2 and 3 weeks old (Illes, Fleischer, et al. 2007). A small subset of electrodes in both culture types also exhibited spontaneous random firing of single action potentials interspersed amongst the bursting activity.

**Figure 2.**
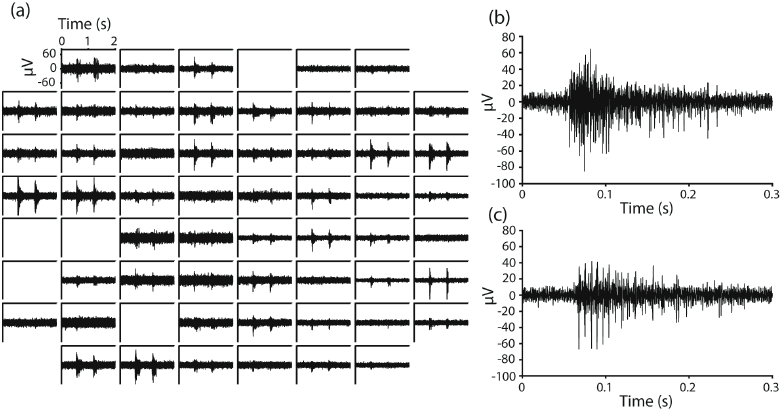
Both dissociated and aggregate cultures display synchronized bursting across MEAs. **(a)** 60 channel electrode arrangement from a single dissociated culture. Each square window represents the extracellular activity recorded from a single electrode over a 2 second period. The channel windows are arranged in the same configuration as the physical electrode contacts on the planar surface of the MEA. Empty windows signify noisy recordings that were set to ground in order to avoid noise interference with neighboring electrodes. **(b)** An enlarged display of the first neuronal burst in the activity recorded on electrode 83 (column 8, row 3). Each individual large spike represents a single action potential. **(c)** An enlarged display of the first neuronal burst in the activity recorded on electrode 38 (column 3, row 8). Each individual large spike represents a single action potential.

### Identification of Individual Neurons from Spontaneous Multi-Unit Activity

Spontaneous MEA spiking activity was thresholded during recordings with a 5RMS threshold to extract the spike waveforms and spike-sorted (**Figure 3a**). The majority of the spike-sorted data exhibited Poisson-like spiking when the inter-spike intervals (ISI) were plotted, which is in concordance with ISI behavior in a variety of previous reports (Perkel, Gerstein, et al. 1967; Longtin, Bulsara, et al. 1991; Jones 2004). **Figure 3b** shows ISI histograms of the same two neurons spikes-sorted from the same electrode. Both histograms are approximately exponentially distributed, implying Poisson-like behavior. **Figure 3c** is a raster plot of 2500 milliseconds of spike-sorted activity from the same two neurons, demonstrating that the differing action potential shapes between the two sorted units was likely not a result of waveform shape changes during extended bursting of a single neuron, which has been previously observed.

**Figure 3.**
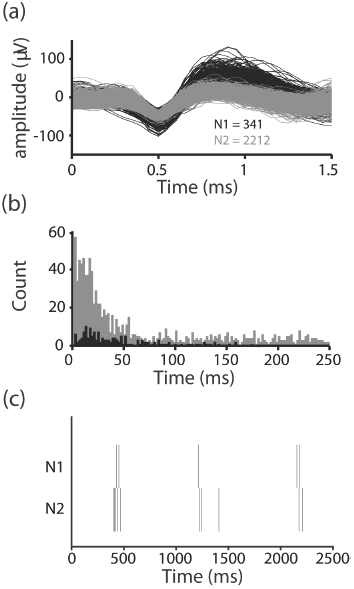
Individual neurons were isolated from single electrodes using offline PCA spike-sorting. **(a)** Two spike-sorted neurons recorded over a 1 hour period from the same electrode in an aggregated culture **(b)** Inter-spike intervals of the two neurons. The black and grey histograms correspond to the black and grey action potential clusters, respectively **(c)** Raster plots of the two neurons over a 2500 millisecond period.

In total, 177 neurons were isolated across 6 dissociated MEA cultures and 162 neurons were isolated across 4 aggregated cultures. **Table 1** provides details about various neuron statistics from these spike-sorted units. One result that arises from the data is the difference in both the absolute number of active units per MEA between the two cultures groups as well as the difference in variability between the groups. Dissociated cultures averaged 19.7 active electrodes (electrodes with at least one neuron having over 100 action potentials during the recording period) out of a total of 64 possible, with a standard deviation of 12.5. On the other hand, aggregated cultures averaged 32 active electrodes with a much smaller standard deviation of 4.7. The numbers of active neurons showed a similar but weaker trend with respective neuron counts of 29.5 ± 19.8 for dissociated and 40.5 ± 9.03 for aggregated. Variability was also low in the spike rate of neurons recorded in aggregated cultures relative to dissociated cultures. All of these neuron data suggest that the aggregate culture protocol may yield more reliable spontaneous spiking in ES-derived networks.

**Table 1.** Culture techniques drive different reliabilities in expected activity. Neurons were determined to be active if they had >100 action potentials over the hour recording period. Electrodes were signified as active if at least one active neuron was recorded from the electrode. Mean spike rate for a single MEA culture was calculated by averaging the spike rate of all active neurons within the culture.

| Culture Type | MEA Culture Number | Num. Active Electrodes | Num. Active Neurons | Mean Spike Rate (sp/s) | Burstiness |
| --- | --- | --- | --- | --- | --- |
| Dissociated Cultures | 1 | 25 | 37 | 1.3 | 0.11 |
|  | 2 | 5 | 9 | 0.56 | 0.49 |
|  | 3 | 29 | 48 | 0.32 | 0.60 |
|  | 4 | 12 | 12 | 0.11 | 0.99 |
|  | 5 | 10 | 16 | 0.28 | 0.72 |
|  | 6 | 37 | 55 | 0.31 | 0.67 |
| | Mean ± STD | 19.7 ± 12.5 | 29.5 ± 19.8 | 0.47 ± 0.41 | 0.60 ± 0.29 |
| Aggregated Cultures | 1 | 33 | 49 | 0.23 | 0.73 |
|  | 2 | 38 | 47 | 0.32 | 0.52 |
|  | 3 | 30 | 36 | 0.30 | 0.57 |
|  | 4 | 27 | 30 | 0.29 | 0.49 |
| | Mean ± STD | 32 ± 4.7 | 40.5 ± 9.03 | 0.28 ± 0.042 | 0.58 ± 0.11 |

Although the distributions of isolated neurons may have varied across culture conditions, the isolated units appeared to maintain similarity in the structure of their firing patterns (**Figure 4**). In **Figure 4a**, a dissociated culture exhibits a very rigid on/off bursting structure for 45 out of the 55 neurons while 10 neurons appear to have a more spontaneous random firing structure. An aggregated culture in **Figure 4b** also has a synchronized bursting response, but it appears to be more widespread across all neurons in the culture, with intermittent random firing between bursts on many of the channels. Burstiness, a global measure of the level of bursting in a neuronal culture (Wagenaar, Madhavan, et al. 2005), is nearly identical between the two groups as shown in **Table 1**. Both culture techniques yield enough neurons after spike-sorting to proceed to an assessment of connectivity in both spatial and temporal dimensions.

**Figure 4.**
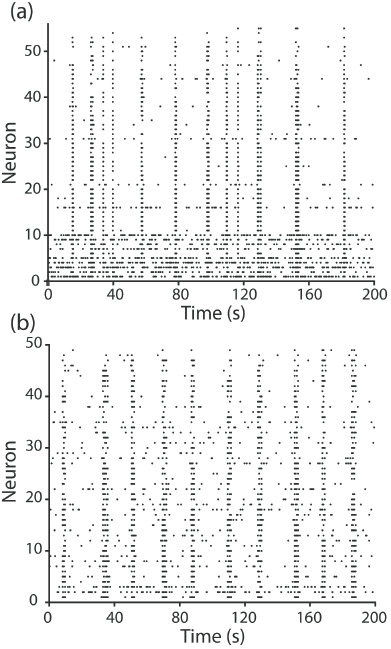
Individual neurons from MEA recordings retain synchronization observed in raw data for both types of cultures. **(a)** Raster plot of action potentials from all of the individual neurons (n = 55) in a single dissociated culture. Each row represents the activity of an individual neuron, where each dot represents a single action potential. Rows were sorted by the number of action potentials in the time period. **(b)** Raster plot of action potentials from all of the individual neurons (n = 49) in a single aggregated culture.

### Cross-Correlation Histograms (CCHs) Represent Neuron-Neuron Functional Connectivity

Using the spike trains of these individual neurons, we then approximated the functional connectivity between individual neurons by calculating each neuron pair’s CCH. This analysis estimates the probability of neuron B firing an action potential at times relative to neuron A’s action potential. Essentially, these values reflect the probabilities over an entire recording period of two neurons firing at each particular time delay relative to one another. The width of the bins used in CCH can be adjusted, and in our particular application, we chose a 5 ms window to maximize the amount of neuron-level correlations captured in the analysis while minimizing network state or noise correlations that occur on broader time scales (Doiron, Litwin-Kumar, et al. 2016). We deemed the resulting histogram the “functional connection” between a neuron pair (**Figure 5**). Typically, peaks within the CCH signify a time relationship between neurons while a flat histogram conveys no relatedness (i.e. a uniform distribution of coincident firing probability).

**Figure 5.**
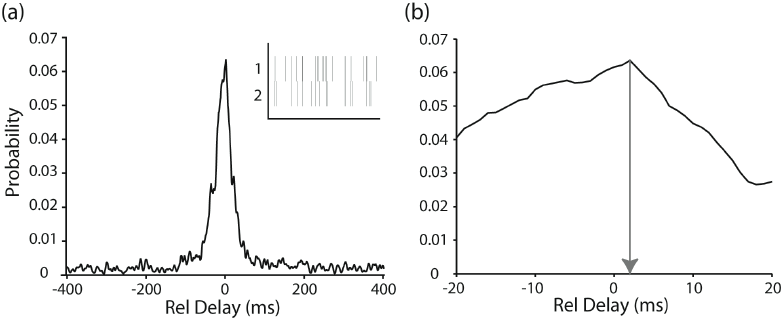
Cross-correlation histograms demonstrate probability of relative spiking across time. **(a)** CCH computed from two different pairs of neurons in dissociated cultures. Inset for each figure are the raster plots of the two neurons being compared, where each row represents the activity from a single neuron for a 40 s period and the dashed lines represent action potentials (b) The peak of the same CCH is located at a time delay of 2 ms.

### Relationship Between Connection Delay and Inter-Neuron Distance in Aggregated Cultures

In order to determine the statistical significance of putative connections between neurons, we chose a methodology that leverages the physical aspects of MEAs and has been previously validated with dissociated mouse suprachiasmatic nucleus (Freeman, Krock, et al. 2013). Determining the significance of connections requires the construction of a null distribution that represents the distribution from which non-connections (CCHs between neurons located in different dishes) are drawn and deriving the significance level at which we can claim with statistical confidence that a connection was not drawn from the null distribution.

**Figures 6 and 7** demonstrate the application of the BSAC methodology for both dissociated (**Figure 6**) and aggregated (**Figure 7**) cultures. A z-score for every CCH was constructed as described in the Materials & Methods to allow for comparisons across CCHs. By comparing neurons that were grown in separate MEAs, we were able to build a null distribution of 879 non-connections for dissociated cultures (**Figure 6a**) and 1079 CCH comparisons in aggregated cultures (**Figure 7a**). As expected, these null distributions were uniform across the 1000 ms delay window, producing the diagonal cumulative curves shown in the upper left panels of **Figure 6a** and **Figure 7a**. The significance values at a 5% false-discovery rate, 13.1 for dissociated and 17.1 for aggregated cultures, were applied to the within-culture z-score distributions for dissociated (**Figure 6b**) and aggregated (**Figure 7b**) cultures, respectively, identifying statistically significant connections lying above the thresholds. For the dissociated and aggregated ES-derived cultures, 837 and 1458 significant connections were detected, and all of the peaks of the significant connections occurred below relative delays of 66 and 61 ms, respectively. These delays are consistent with the timing that might be expected for network interactions.

**Figure 6.**
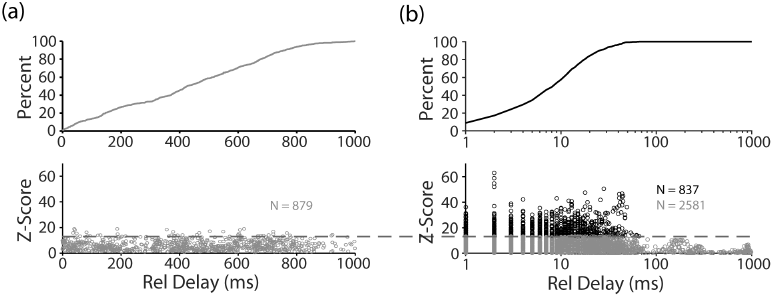
BSAC methodology detects statistically significant neuron-neuron functional connections in dissociated cultures. Black and grey circles represent the z-score of a single cross-correlogram. **(a)** Bottom-panel represents the z-score distribution of cross-correlations computed between neurons located in separate dissociated MEA cultures. These “virtual” non-connections were computed to serve as the chance distribution for calculating significance. A false-discovery rate of 0.05 was selected to determine the z-score significance level, 13.1, shown as the dark grey dashed line **(b)** Bottom-panel represents the distribution of cross-correlation z-scores calculated from neurons grown within the same dissociated MEA cultures, compiled across all dissociated cultures (n=6). Altogether, 837 statistically significant connections were detected using the previously calculated significance level (13.1).

**Figure 7.**
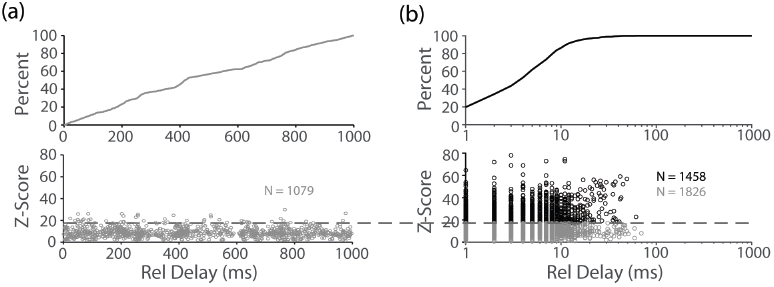
BSAC methodology detects statistically significant neuron-neuron functional connections in aggregated cultures. Black and grey circles represent the z-score of a single cross-correlogram. **(a)** Bottom-panel represents the z-score distribution of cross-correlations computed between neurons located in separate dissociated MEA cultures. A false-discovery rate of 0.05 was selected to determine the z-score significance level, 17.1, shown as the dark grey dashed line **(b)** Bottom-panel represents the distribution of cross-correlation z-scores calculated from neurons grown within the same dissociated MEA cultures, compiled across all dissociated cultures (n=4). Altogether, 1458 statistically significant connections were detected using the previously calculated significance level (17.1).

With significant functional connections identified, the properties of these connections could then be evaluated, in particular, the spatiotemporal nature of the individual neuron-neuron connections. In the context of spinal cord injury repair, ESC-derived graft networks would ideally serve as relays across the site of injury, suggesting the neuronal interconnectivity of a therapeutic candidate population should behave temporally like *in vivo* neurons separated by one or more direct synapse: increasing distance should increase the average relative delay in the spiking of the neuron pair in question. **Figure 8** shows a comparison of this spatiotemporal pattern between dissociated and aggregated ESC-derived cultures. When the functional connection delay between 837 statistically significant neuron pairs in dissociated cultures was grouped by each neuron pair’s corresponding inter-electrode distance, averaged, and plotted against inter-electrode distance, dissociated cultures did not demonstrate a strong relationship between distance and delay (**Figure 8a**). Repeating this analysis for the 1458 statistically significant neuron pairs in aggregated cultures, however, revealed a very tight linear relationship between distance and connection delay for distances below 1200 μm (**Figure 8b**), suggesting that synaptically connected neurons within aggregated cultures tended to have increasing relative delays in the timing of action potentials as the distance between connected neurons increased.

**Figure 8.**
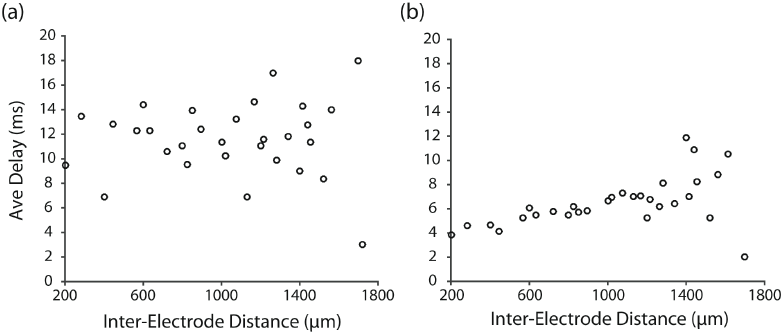
Neuron-Neuron Distance Related to Connection Delay. (a) Functional connections within dissociated cultures were grouped according to the distance between the two electrodes from which the two neurons in each connection were recorded. Inter-electrode distances exist in discrete bins due to the nature of MEAs, where adjacent electrodes in the MEA dishes were located 200 μm apart in an 8x8 grid. Neurons were assumed to be located at the electrode at which their respective action potentials were identified. The mean connection delay for each inter-electrode distance bin was taken and plotted against inter-electrode distance (b) The previous analysis was repeated for aggregated cultures.

## DISCUSSION

To our knowledge, this study represents the first comparison of the individual neuron-neuron functional connections in aggregate, or neurosphere, MEA culture relative to traditional dissociated culture for investigating ESC-derived neuronal interconnectivity. The 3D-like structure of aggregate culture provides a favorable growth-promoting environment for neuronal and glial cells, thus leading to more live-tissue like properties (Pampaloni, Reynaud, et al. 2007; Lu, Searle, et al. 2012; Edmondson, Broglie, et al. 2014). Recently, aggregate cultures have been extensively used to characterize global network activity in ESC-derived MEA cultures (Matthieu, Honegger, et al. 1978; Heikkila, Yla-Outinen, et al. 2009; Koito and Li 2009; Lappalainen, Salomaki, et al. 2010; Illes, Jakab, et al. 2014). Of these studies, only one demonstrated the effects of neural aggregation on network activity, showing that aggregation led to an increase in synchronous bursting activity across electrodes relative to dissociated cultures (Illes, Theiss, et al. 2009).

Our data suggest that both culture types can support spontaneously active Chx10-enriched cultures with bursting behavior reminiscent of previous ESC-derived MEA investigations (Ban, Bonifazi, et al. 2007; Illes, Fleischer, et al. 2007; Heikkila, Yla-Outinen, et al. 2009; Lappalainen, Salomaki, et al. 2010; Illes, Jakab, et al. 2014). Interestingly, both dissociated and aggregated Chx10-enriched cultures had nearly identical levels of bursting across the recording period as measured by the “burstiness” metric presented and utilized by Wagenaar and colleagues in multiple investigations (Wagenaar, Madhavan, et al. 2005; Wagenaar, Pine, et al. 2006; Wagenaar, Pine, et al. 2006), but only aggregate Chx10-enriched cultures support repeatable electrode coverage and neuron detection compared to the high variability in dissociated MEA cultures. The synchronized bursting activity observed in both cultures conditions is also consistent with the ESC-derived activity characterized in previous studies (Ban, Bonifazi, et al.; Illes, Fleischer, et al.; Heikkila, Yla-Outinen, et al.; Lappalainen, Salomaki, et al.; Illes, Jakab, et al.). Global bursts are ubiquitous across both primary and ESC-derived cultures, but the bursting in our spinal cord ESC-derived cultures occurred at a slower burst rate compared to that of primary cultures (Maeda, Robinson, et al. 1995; Wagenaar, Madhavan, et al. 2005; Wagenaar, Pine, et al. 2006).

Cross-correlation histograms have been ubiquitously applied in the neuronal assembly literature to characterize the temporal relatedness between simultaneously recorded neurons by quantitatively and qualitatively describing neuronal connections with direct synaptic interactions (Herrmann and Gerstner 2002; Bartho, Hirase, et al. 2004; Veredas, Vico, et al. 2005; Fujisawa, Amarasingham, et al. 2008; Freeman, Krock, et al. 2013), common input (Kirkwood and Sears 1978; Binder and Powers 2001; Turker and Powers 2002), and background network activity effects (Aertsen, Gerstein, et al. 1989; Constantinidis, Franowicz, et al. 2001; Ostojic, Brunel, et al. 2009). The width of the time bins used in the calculations can also influence the type of connectivity encompassed by the histograms. State-dependent correlations in neural activity occur at longer time-scales (Doiron, Litwin-Kumar, et al. 2016). ESC-derived MEA studies that utilize wider binning in their connectivity measures can lead to overestimates in neuron-neuron correlations (Ban, Bonifazi, et al. 2007).

Typically, these measurements are interpreted as to identify the physical neural circuitry underlying the measured connectivity. For example, thin peaks can signify the existence of monosynaptic connections or shared excitatory input (Perkel, Gerstein, et al. 1967; Fujisawa, Amarasingham, et al. 2008; Freeman, Krock, et al. 2013), while broad peaks may represent shared inhibitory synaptic input (Perkel, Gerstein, et al. 1967; Perkel, Gerstein, et al. 1967; Aertsen, Gerstein, et al. 1989). Any interpretation of circuitry requires 1) the assumption of stationarity, meaning the spike rates of the simultaneously recorded neurons remain stable over various time scales (Moore, Perkel, et al. 1966; Perkel, Gerstein, et al. 1967; Perkel, Gerstein, et al. 1967; Aertsen, Gerstein, et al. 1989), and 2) an inspection of the auto-correlations of the individual neurons themselves (Moore, Perkel, et al. 1966; Perkel, Gerstein, et al. 1967). Due to the prominent synchronized bursting and long periods of silence in our recorded data, we cannot make a stationarity assumption, which limits what can be ascertained about the circuitry underlying each individual connection.

This outcome does not, however, diminish the utility of these statistically significant functional connections for connectivity comparisons in the context of diagnostics. Resting state functional connectivity has become an extremely useful diagnostic tool for cortical disorders even though the circuit diagrams underlying the observed correlations between cortical areas are not well defined (Fox and Greicius 2010; Woodward and Cascio 2015; Ann, Jun, et al. 2016). Similarly, neuron-neuron functional connectivity can be used as a means to assess differences in the functional properties of ESC-derived neuron cultures across culture-type. As shown in the present study, MEAs provide a means to rapidly identify thousands of individual connections and characterize how distributions of these connections change across groups. Although Chx10-enriched cultures in both dissociated and aggregated groups consistently contained hundreds of identifiable functional connections with the use of CCHs and BSAC, aggregated cultures not only contained a larger quantity of significant connections across fewer cultures, but also demonstrated a positive tight linear relationship between neuron distance and connection delay for short-range (<1200 μm) connections.

As stated previously, consistent and predictable signal propagation is desirable for any ESC-derived population that is to be used as a grafted relay across the site of spinal cord injury (Bonner, Connors, et al. 2011; Lu, Wang, et al. 2012; Bonner and Steward 2015). Our results suggest that neural aggregation of ESC-derived cultures supports this type of consistent spatiotemporal relationship in neuron-neuron functional connectivity as compared to dissociated cultures. This finding may limit future investigations to aggregated cultures so as to focus on characteristic differences between ESC-derived cultures of different genetically-defined neuronal populations. This MEA-based assay could also be extended to construct adjacency matrices for network topological comparisons across populations, but shortcomings of current mathematical frameworks likely need to be overcome to perform such an analysis (van Wijk, Stam, et al. 2010).

## CONCLUSION

We have shown that MEAs, stem cell differentiation protocols, and computational techniques can be combined to assess network connectivity of ESC-derived neuronal populations. Our ability to leverage the nature of MEA cultures to calculate connection significance not only further validates a methodology pioneered in other systems neuroscience applications (Freeman, Krock, et al. 2013), but also demonstrates its promise in characterizing the connectivity of ESC-derived transplantation candidates. Such methodologies can be expanded to explore the neurotransmitters underlying neuronal connections and the differential effects of stimulation on ESC-derived neuronal networks. Moving forward, these types of approaches, in parallel with *in vivo* transplantation studies in animal models of SCI, will provide an avenue towards generating network fingerprints of therapeutically successful ESC-relay grafting.

## ACKNOWLEDGEMENTS

The authors acknowledge funding from the following sources: National Institute of Neurological Disorders and Stroke of the National Institutes of Health under grant R01 NS090617, Individual Predoctoral National Research Service Award (F31NS090760), and the University of Missouri System Spinal Cord Injuries Research Program.

## REFERENCES

Abematsu, M., Tsujimura, K., Yamano, M., Saito, M., Kohno, K., Kohyama, J., Namihira, M., Komiya, S. and Nakashima, K. (2010). “Neurons derived from transplanted neural stem cells restore disrupted neuronal circuitry in a mouse model of spinal cord injury.” J Clin Invest 120(9): 3255–3266.

Aertsen, A., Vaadia, E., Abeles, M., Ahissar, E., Bergman, H., Karmon, B., Lavner, Y., Margalit, E., Nelken, I. and Rotter, S. (1991). “Neural interactions in the frontal cortex of a behaving monkey: signs of dependence on stimulus context and behavioral state.” J Hirnforsch 32(6): 735–743.

Aertsen, A. M., Gerstein, G. L., Habib, M. K. and Palm, G. (1989). “Dynamics of neuronal firing correlation: modulation of “effective connectivity”.” J Neurophysiol 61(5): 900–917.

Anderson, D., Self, T., Mellor, I. R., Goh, G., Hill, S. J. and Denning, C. (2007). “Transgenic enrichment of cardiomyocytes from human embryonic stem cells.” Mol Ther 15(11): 2027–2036.

Ann, H. W., Jun, S., Shin, N. Y., Han, S., Ahn, J. Y., Ahn, M. Y., Jeon, Y. D., Jung, I. Y., Kim, M. H., Jeong, W. Y., Ku, N. S., Kim, J. M., Smith, D. M. and Choi, J. Y. (2016). “Characteristics of Resting-State Functional Connectivity in HIV-Associated Neurocognitive Disorder.” PLoS One 11(4): e0153493.

Ban, J., Bonifazi, P., Pinato, G., Broccard, F. D., Studer, L., Torre, V. and Ruaro, M. E. (2007). “Embryonic stem cell-derived neurons form functional networks in vitro.” Stem Cells 25(3): 738–749.

Bartho, P., Hirase, H., Monconduit, L., Zugaro, M., Harris, K. D. and Buzsaki, G. (2004). “Characterization of neocortical principal cells and interneurons by network interactions and extracellular features.” J Neurophysiol 92(1): 600–608.

Binder, M. D. and Powers, R. K. (2001). “Relationship between simulated common synaptic input and discharge synchrony in cat spinal motoneurons.” J Neurophysiol 86(5): 2266–2275.

Boehler, M. D., Leondopulos, S. S., Wheeler, B. C. and Brewer, G. J. (2012). “Hippocampal networks on reliable patterned substrates.” J Neurosci Methods 203(2): 344–353.

Bonner, J. F., Connors, T. M., Silverman, W. F., Kowalski, D. P., Lemay, M. A. and Fischer, I. (2011). “Grafted neural progenitors integrate and restore synaptic connectivity across the injured spinal cord.” J Neurosci 31(12): 4675–4686.

Bonner, J. F. and Steward, O. (2015). “Repair of spinal cord injury with neuronal relays: From fetal grafts to neural stem cells.” Brain Res 1619: 115–123.

Brown, C. R., Butts, J. C., McCreedy, D. A. and Sakiyama-Elbert, S. E. (2014). “Generation of v2a interneurons from mouse embryonic stem cells.” Stem Cells Dev 23(15): 1765–1776.

Brustle, O., Jones, K. N., Learish, R. D., Karram, K., Choudhary, K., Wiestler, O. D., Duncan, I. D. and McKay, R. D. (1999). “Embryonic stem cell-derived glial precursors: a source of myelinating transplants.” Science 285(5428): 754–756.

Chao, Z. C., Bakkum, D. J. and Potter, S. M. (2007). “Region-specific network plasticity in simulated and living cortical networks: comparison of the center of activity trajectory (CAT) with other statistics.” J Neural Eng 4(3): 294–308.

Constantinidis, C., Franowicz, M. N. and Goldman-Rakic, P. S. (2001). “Coding specificity in cortical microcircuits: a multiple-electrode analysis of primate prefrontal cortex.” J Neurosci 21(10): 3646–3655.

Cutts, C. S. and Eglen, S. J. (2014). “Detecting pairwise correlations in spike trains: an objective comparison of methods and application to the study of retinal waves.” J Neurosci 34(43): 14288–14303.

Doiron, B., Litwin-Kumar, A., Rosenbaum, R., Ocker, G. K. and Josic, K. (2016). “The mechanics of state-dependent neural correlations.” Nat Neurosci 19(3): 383–393.

Downes, J. H., Hammond, M. W., Xydas, D., Spencer, M. C., Becerra, V. M., Warwick, K., Whalley, B. J. and Nasuto, S. J. (2012). “Emergence of a small-world functional network in cultured neurons.” PLoS Comput Biol 8(5): e1002522.

Duncan, I. D., Aguayo, A. J., Bunge, R. P. and Wood, P. M. (1981). “Transplantation of rat Schwann cells grown in tissue culture into the mouse spinal cord.” J Neurol Sci 49(2): 241–252.

Edmondson, R., Broglie, J. J., Adcock, A. F. and Yang, L. (2014). “Three-dimensional cell culture systems and their applications in drug discovery and cell-based biosensors.” Assay Drug Dev Technol 12(4): 207–218.

Fox, M. D. and Greicius, M. (2010). “Clinical applications of resting state functional connectivity.” Front Syst Neurosci 4: 19.

Freeman, G. M., Jr., Krock, R. M., Aton, S. J., Thaben, P. and Herzog, E. D. (2013). “GABA networks destabilize genetic oscillations in the circadian pacemaker.” Neuron 78(5): 799–806.

Friston, K. J. (1994). “Functional and Effective Connectivity in Neuroimaging: A Synthesis.” Human Brain Mapping(2): 56–78.

Friston, K. J. (2011). “Functional and effective connectivity: a review.” Brain Connect 1(1): 13–36.

Friston, K. J., Frith, C. D., Liddle, P. F. and Frackowiak, R. S. (1993). “Functional connectivity: the principal-component analysis of large (PET) data sets.” J Cereb Blood Flow Metab 13(1): 5–14.

Fujimoto, Y., Abematsu, M., Falk, A., Tsujimura, K., Sanosaka, T., Juliandi, B., Semi, K., Namihira, M., Komiya, S., Smith, A. and Nakashima, K. (2012). “Treatment of a mouse model of spinal cord injury by transplantation of human induced pluripotent stem cell-derived long-term self-renewing neuroepithelial-like stem cells.” Stem Cells 30(6): 1163–1173.

Fujisawa, S., Amarasingham, A., Harrison, M. T. and Buzsaki, G. (2008). “Behavior-dependent short-term assembly dynamics in the medial prefrontal cortex.” Nat Neurosci 11(7): 823–833.

Garofalo, M., Nieus, T., Massobrio, P. and Martinoia, S. (2009). “Evaluation of the performance of information theory-based methods and cross-correlation to estimate the functional connectivity in cortical networks.” PLoS One 4(8): e6482.

Gerstein, G. L. and Perkel, D. H. (1972). “Mutual temporal relationships among neuronal spike trains. Statistical techniques for display and analysis.” Biophys J 12(5): 453–473.

Heikkila, T. J., Yla-Outinen, L., Tanskanen, J. M., Lappalainen, R. S., Skottman, H., Suuronen, R., Mikkonen, J. E., Hyttinen, J. A. and Narkilahti, S. (2009). “Human embryonic stem cell-derived neuronal cells form spontaneously active neuronal networks in vitro.” Exp Neurol 218(1): 109–116.

Herrmann, A. and Gerstner, W. (2002). “Noise and the PSTH response to current transients: II. Integrate-and-fire model with slow recovery and application to motoneuron data.” J Comput Neurosci 12(2): 83–95.

Hou, S., Tom, V. J., Graham, L., Lu, P. and Blesch, A. (2013). “Partial restoration of cardiovascular function by embryonic neural stem cell grafts after complete spinal cord transection.” J Neurosci 33(43): 17138–17149.

Illes, S., Fleischer, W., Siebler, M., Hartung, H. P. and Dihne, M. (2007). “Development and pharmacological modulation of embryonic stem cell-derived neuronal network activity.” Exp Neurol 207(1): 171–176.

Illes, S., Jakab, M., Beyer, F., Gelfert, R., Couillard-Despres, S., Schnitzler, A., Ritter, M. and Aigner, L. (2014). “Intrinsically active and pacemaker neurons in pluripotent stem cell-derived neuronal populations.” Stem Cell Reports 2(3): 323–336.

Illes, S., Theiss, S., Hartung, H. P., Siebler, M. and Dihne, M. (2009). “Niche-dependent development of functional neuronal networks from embryonic stem cell-derived neural populations.” BMC Neurosci 10: 93.

Iyer, N. R., Huettner, J. E., Butts, J. C., Brown, C. R. and Sakiyama-Elbert, S. E. (2016). “Generation of highly enriched V2a interneurons from mouse embryonic stem cells.” Exp Neurol 277: 305–316.

Jones, B. E. (2004). “Activity, modulation and role of basal forebrain cholinergic neurons innervating the cerebral cortex.” Prog Brain Res 145: 157–169.

Kirkwood, P. A. and Sears, T. A. (1978). “The synaptic connexions to intercostal motoneurones as revealed by the average common excitation potential.” J Physiol 275: 103–134.

Koito, H. and Li, J. (2009). “Preparation of rat brain aggregate cultures for neuron and glia development studies.” J Vis Exp(31).

Lappalainen, R. S., Salomaki, M., Yla-Outinen, L., Heikkila, T. J., Hyttinen, J. A., Pihlajamaki, H., Suuronen, R., Skottman, H. and Narkilahti, S. (2010). “Similarly derived and cultured hESC lines show variation in their developmental potential towards neuronal cells in long-term culture.” Regen Med 5(5): 749–762.

Lee, K. Z., Lane, M. A., Dougherty, B. J., Mercier, L. M., Sandhu, M. S., Sanchez, J. C., Reier, P. J. and Fuller, D. D. (2014). “Intraspinal transplantation and modulation of donor neuron electrophysiological activity.” Exp Neurol 251: 47–57.

Li, M., Pevny, L., Lovell-Badge, R. and Smith, A. (1998). “Generation of purified neural precursors from embryonic stem cells by lineage selection.” Curr Biol 8(17): 971–974.

Longtin, A., Bulsara, A. and Moss, F. (1991). “Time-interval sequences in bistable systems and the noise-induced transmission of information by sensory neurons.” Phys Rev Lett 67(5): 656–659.

Lu, H., Searle, K., Liu, Y. and Parker, T. (2012). “The effect of dimensionality on growth and differentiation of neural progenitors from different regions of fetal rat brain in vitro: 3-dimensional spheroid versus 2-dimensional monolayer culture.” Cells Tissues Organs 196(1): 48–55.

Lu, P., Wang, Y., Graham, L., McHale, K., Gao, M., Wu, D., Brock, J., Blesch, A., Rosenzweig, E. S., Havton, L. A., Zheng, B., Conner, J. M., Marsala, M. and Tuszynski, M. H. (2012). “Long-distance growth and connectivity of neural stem cells after severe spinal cord injury.” Cell 150(6): 1264–1273.

Maccione, A., Gandolfo, M., Tedesco, M., Nieus, T., Imfeld, K., Martinoia, S. and Berdondini, L. (2010). “Experimental Investigation on Spontaneously Active Hippocampal Cultures Recorded by Means of High-Density MEAs: Analysis of the Spatial Resolution Effects.” Front Neuroeng 3: 4.

Maccione, A., Garofalo, M., Nieus, T., Tedesco, M., Berdondini, L. and Martinoia, S. (2012). “Multiscale functional connectivity estimation on low-density neuronal cultures recorded by high-density CMOS Micro Electrode Arrays.” J Neurosci Methods 207(2): 161–171.

Maeda, E., Robinson, H. P. and Kawana, A. (1995). “The mechanisms of generation and propagation of synchronized bursting in developing networks of cortical neurons.” J Neurosci 15(10): 6834–6845.

Marconi, E., Nieus, T., Maccione, A., Valente, P., Simi, A., Messa, M., Dante, S., Baldelli, P., Berdondini, L. and Benfenati, F. (2012). “Emergent functional properties of neuronal networks with controlled topology.” PLoS One 7(4): e34648.

Matthieu, J. M., Honegger, P., Trapp, B. D., Cohen, S. R. and Webster, H. F. (1978). “Myelination in rat brain aggregating cell cultures.” Neuroscience 3(6): 565–572.

McCreedy, D. A., Brown, C. R., Butts, J. C., Xu, H., Huettner, J. E. and Sakiyama-Elbert, S. E. (2014). “A New Method for Generating High Purity Motoneurons From Mouse Embryonic Stem Cells.” Biotechnol Bioeng 111(10): 2041–2055.

McCreedy, D. A., Rieger, C. R., Gottlieb, D. I. and Sakiyama-Elbert, S. E. (2012). “Transgenic enrichment of mouse embryonic stem cell-derived progenitor motor neurons.” Stem Cell Res 8(3): 368–378.

McCreedy, D. A., Wilems, T. S., Xu, H., Butts, J. C., Brown, C. R., Smith, A. W. and Sakiyama-Elbert, S. E. (2014). “Survival, Differentiation, and Migration of High-Purity Mouse Embryonic Stem Cell-derived Progenitor Motor Neurons in Fibrin Scaffolds after Sub-Acute Spinal Cord Injury.” Biomater Sci 2(11): 1672–1682.

Moore, G. P., Perkel, D. H. and Segundo, J. P. (1966). “Statistical analysis and functional interpretation of neuronal spike data.” Annu Rev Physiol 28: 493–522.

Mothe, A. J., Tam, R. Y., Zahir, T., Tator, C. H. and Shoichet, M. S. (2013). “Repair of the injured spinal cord by transplantation of neural stem cells in a hyaluronan-based hydrogel.” Biomaterials 34(15): 3775–3783.

Nori, S., Okada, Y., Yasuda, A., Tsuji, O., Takahashi, Y., Kobayashi, Y., Fujiyoshi, K., Koike, M., Uchiyama, Y., Ikeda, E., Toyama, Y., Yamanaka, S., Nakamura, M. and Okano, H. (2011). “Grafted human-induced pluripotent stem-cell-derived neurospheres promote motor functional recovery after spinal cord injury in mice.” Proc Natl Acad Sci U S A 108(40): 16825–16830.

Ostojic, S., Brunel, N. and Hakim, V. (2009). “How connectivity, background activity, and synaptic properties shape the cross-correlation between spike trains.” J Neurosci 29(33): 10234–10253.

Pampaloni, F., Reynaud, E. G. and Stelzer, E. H. (2007). “The third dimension bridges the gap between cell culture and live tissue.” Nat Rev Mol Cell Biol 8(10): 839–845.

Pastore, V. P., Poli, D., Godjoski, A., Martinoia, S. and Massobrio, P. (2016). “ToolConnect: A Functional Connectivity Toolbox for In vitro Networks.” Front Neuroinform 10: 13.

Perkel, D. H., Gerstein, G. L. and Moore, G. P. (1967). “Neuronal spike trains and stochastic point processes. I. The single spike train.” Biophys J 7(4): 391–418.

Perkel, D. H., Gerstein, G. L. and Moore, G. P. (1967). “Neuronal spike trains and stochastic point processes. II. Simultaneous spike trains.” Biophys J 7(4): 419–440.

Poli, D., Pastore, V. P. and Massobrio, P. (2015). “Functional connectivity in in vitro neuronal assemblies.” Front Neural Circuits 9: 57.

Schroeter, M. S., Charlesworth, P., Kitzbichler, M. G., Paulsen, O. and Bullmore, E. T. (2015). “Emergence of rich-club topology and coordinated dynamics in development of hippocampal functional networks in vitro.” J Neurosci 35(14): 5459–5470.

Sharp, K. G., Yee, K. M. and Steward, O. (2014). “A re-assessment of long distance growth and connectivity of neural stem cells after severe spinal cord injury.” Exp Neurol 257: 186–204.

Tetzlaff, W., Okon, E. B., Karimi-Abdolrezaee, S., Hill, C. E., Sparling, J. S., Plemel, J. R., Plunet, W. T., Tsai, E. C., Baptiste, D., Smithson, L. J., Kawaja, M. D., Fehlings, M. G. and Kwon, B. K. (2011). “A systematic review of cellular transplantation therapies for spinal cord injury.” J Neurotrauma 28(8): 1611–1682.

Turker, K. S. and Powers, R. K. (2002). “The effects of common input characteristics and discharge rate on synchronization in rat hypoglossal motoneurones.” J Physiol 541(Pt 1): 245–260.

van Wijk, B. C., Stam, C. J. and Daffertshofer, A. (2010). “Comparing brain networks of different size and connectivity density using graph theory.” PLoS One 5(10): e13701.

Veredas, F. J., Vico, F. J. and Alonso, J. M. (2005). “Factors determining the precision of the correlated firing generated by a monosynaptic connection in the cat visual pathway.” J Physiol 567(Pt 3): 1057–1078.

Wagenaar, D. A., Madhavan, R., Pine, J. and Potter, S. M. (2005). “Controlling bursting in cortical cultures with closed-loop multi-electrode stimulation.” J Neurosci 25(3): 680–688.

Wagenaar, D. A., Pine, J. and Potter, S. M. (2006). “An extremely rich repertoire of bursting patterns during the development of cortical cultures.” BMC Neurosci 7: 11.

Wagenaar, D. A., Pine, J. and Potter, S. M. (2006). “Searching for plasticity in dissociated cortical cultures on multi-electrode arrays.” J Negat Results Biomed 5: 16.

Woodward, N. D. and Cascio, C. J. (2015). “Resting-State Functional Connectivity in Psychiatric Disorders.” JAMA Psychiatry 72(8): 743–744.

Xu, H., Iyer, N., Huettner, J. E. and Sakiyama-Elbert, S. E. (2015). “A puromycin selectable cell line for the enrichment of mouse embryonic stem cell-derived V3 interneurons.” Stem Cell Res Ther 6: 220.

Xu, H. and Sakiyama-Elbert, S. E. (2015). “Directed Differentiation of V3 Interneurons from Mouse Embryonic Stem Cells.” Stem Cells Dev 24(22): 2723–2732.

